# Identifying key species in meta-communities

**DOI:** 10.1101/2023.09.14.557694

**Authors:** Guillaume Rollin, José Lages, Benoit Gauzens

**Affiliations:** Équipe de physique théorique, Institut UTINAM, CNRS, Université de Franche-Comté, Besançon, France; EcoNetLab, German Centre for Integrative Biodiversity Research (iDiv) Halle-Jena-Leipzig, Leipzig, Germany; Institute of Biodiversity, Friedrich Schiller University Jena 07743 Germany

## Abstract

1. With the ongoing biodiversity crisis, identifying which species are of particular importance to prevent the extinction of other species has become a pressing issue. However, most approaches to detect these important species are made at a local (i.e, community) level, without considering the potential effect of species dispersion in a landscape.
2. We present a modified PageRank algorithm to determine the importance of species in meta-communities based on two sets of networks: food webs that depict local trophic interactions and landscape networks representing the movement of species across different habitat patches.
3. We show that (i) what is considered an important species changes between isolated communities and meta-communities and (ii) the importance of a species in a meta-community depends on the position of its habitat patch in the landscape network.
4. Our results stress the need for a global consideration of space in the identification of important species.

## Introduction

In the context of the ongoing biodiversity crisis, predicting the effects of species extinctions on other species is a pressing issue. In ecosystems, losing a species can lead to cascading effects on others species because of the complex set of ecological dependencies that species have with each other. For instance, after the loss of a species, some consumers can be left without further access to resources leading them to extinction (secondary extinction). Within a community, food webs compile the set of species and their trophic interactions and can be used to depict species inter-dependencies associated with their need for energy. They thus offer a valuable tool to identify which species are key to maintain the energetic integrity of communities and minimise the number of secondary extinctions Solé and Montoya, 2001]. Therefore, several methods relying on food webs exists to identify important species. First, species focused methods consider information at the species level, defining importance based on the number of interactions or consumers a species has Dunne et al., 2002]. Second, local approaches consider the neighborhood of species: important species are the ones with a high number of connections to other species being themselves connected to a high number of other species, this definition being potentially iterated to consider a more distant neighborhood Jordán, 2009]. Finally, global approaches consider the entire network to define species importance. For instance, Allesina and Pascual Allesina and Pascual, 2009] adapted the Brin and Page PageRank algorithm Brin and Page, 1998] to define species by their ability to spread nutrients (so energy) in the entire food web.

A limitation of these approaches is to consider that ecological communities live in isolation from each other. Indeed, individuals can disperse among landscapes and therefore connect different habitat patches. While food webs depict the possible pathways for energy propagation among species of the same community, landscape networks -that describe how species movement connects different habitat patches in a landscape -depict the possible pathways for energy spreading across the space. The integration of both networks, where food webs are entangled in landscape networks forms meta-food webs that depict transfers of energy in meta-communities (Fig. 1a). This spatial interconnection of different local food webs is associated with mechanisms that can change species persistence locally. For instance, some species that would have gone extinct in a local community can persist because of the constant immigration of individuals from neighbouring habitat patches. This rescue effect Brown and Kodric-Brown, 1977] can occur directly, when individuals of a given species support the local population of the same species Brown and Kodric-Brown, 1977] or indirectly, when this inflow of individuals creates more favourable biotic conditions locally, for instance by increasing the amount of resources available Ryser et al., 2019]. Therefore, the energetic integrity of a local community not only depends on species’ ability to acquire and spread resources through trophic interactions but also on how they are connected to other habitat patches in a landscape. This spatially explicit view of ecological communities is so far ignored by methods aiming at identifying important species in ecosystems and therefore questions our ability to derive sound conservation practices at the landscape level. Indeed, species with a high ability to spread energy within food webs are not necessarily the ones that would spread energy towards different habitat patches, which implies that what is considered an important species in a local community can differ from the real-world context where communities are connected to each other. We here extend the work of Allesina and Pascual Allesina and Pascual, 2009] -that adapted the PageRank algorithm used to sort internet pages by importance to food webs -to propose a method able to identify key species in a meta-community context. We first present how species importance can be estimated for food webs and landscape networks separately before developing the integration of both. We then use our framework to show (1) that species that are considered as important in local communities will change depending on the location of the community in the landscape network and (2) that the location of a patch in the landscape will affect the importance of the species it hosts.

**Figure 1.**
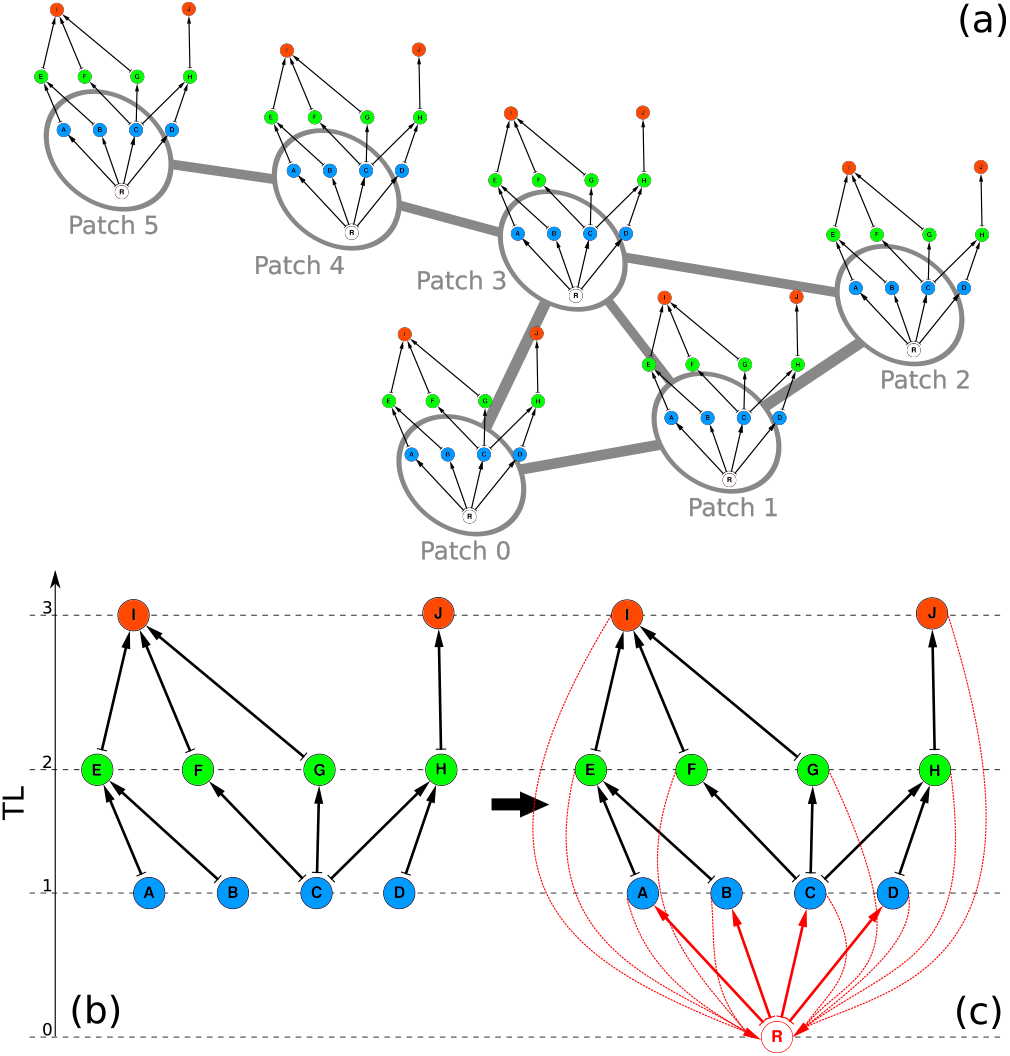
Panel (a) : typical example of meta-food web 𝕄. The underlying landscape network 𝕃 is a non directed network represented here in grey color. To each patch of the landscape, a clone of a rooted food web 𝔽 is attached. The rooted food web is presented below on panel (c). The grey inter-patch links allow the spatial diffusion of species over the landscape. **Panel (b) : typical example of food web**. In blue, A,B,C and D species find their nutrients only in the environment, they are the primary producers and their trophic level (TL) is 1. In green, E,F,G and H species are the predator of the primary producers and their trophic level is 2. Finally, in red, I and J species are the top predators and their trophic level is 3. **Panel (c) : *food web*** 𝔽. The root node, R, is an extra node which represents the non-living nutrients. The red dotted arrows represent the added links which preserve the random walk from the sub-networks problem Langville and Meyer, 2012]. They represent the inevitable species loss of matter which is recycled. The red solid lines indicate that primary producers obtain their nutrients from the environment represented by the root R node. This transformation of a food web into its corresponding rooted food web can be found in Allesina and Pascual, 2009].

## 1 Quantifying importance in different types of networks

### 1.1 PageRank and CheiRank algorithms for food web

The original PageRank algorithm Langville and Meyer, 2012, Brin and Page, 1998] models a random walk on the WWW in order to assess the importance of internet pages. It mimics the journey of a web surfer that follows randomly the succession of hyperlinks leading to new web pages. In this case, the WWW is a complex network whose nodes are the web pages and the directed edges are the hyperlinks. This complex network can be represented by the adjacency matrix *A* in which the presence or absence of a link from page *i* to page *j* is defined by the matrix entry *A*_*ji*_. The PageRank algorithm relies on a stochastic matrix *S* that describes the probabilities of transition between nodes in the network, defined as

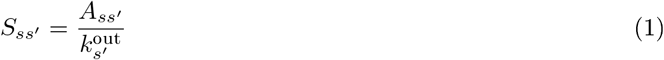

where 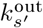 is the out-degree of page *s*′ (i.e. the number of hyperlinks in page *s*′).

This network approach allows for an analogy with food webs, for which the adjacency matrix represents the presence or absence of trophic interaction (the edges) between species (the nodes). The PageRank algorithm therefore can - and was -applied to food webs to estimate species importances Allesina and Pascual, 2009]. However, to use the PageRank algorithm, the stochastic matrix *S* associated with the network has to be irreducible and primitive. Following the solution from Allesina and Pascual, 2009], we ensure these properties in the case of a food web by adding a root node (see Fig. 1c) which can be thought as representing the environment from which primary producers capture energy and towards which all the species can be recycled. To estimate nodes importance, we start by assuming an initial distribution 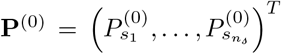. The element 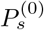, where *s* stands for anyone of the *n*_𝓈_ species, say *s*_1_, *s*_2_, …, *n*_𝓈_, gives the initial probability of presence of a hypothetical random surfer on the node *s*. Hence, after the *m*th iteration the probability of ending up on the site *s*′ is 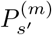 given by **P**^(*m*)^ = ***S***^*m*^**P**^(0)^. The steady state of such a stochastic process is characterized by the PageRank vector 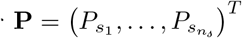 defined such as ***S*P** = **P**. Sorting then the species by descending order of the value of their PageRank probabilities, ie *P*_*s*_ for the species *s*, allows to rank the species in accordance to their ability to catch nutrients from the rooted food web 𝔽. For each species *s* we assign a PageRank index *k*_*s*_ ∈ *{*1, …, *n*_*s*_*}* such as if *P*_*s*_ *> P*_*s*_′, ie if the species *s* is a more efficient nutrient catcher than the species *s*′, then *k*_*s*_ *< k*_*s*_′, and conversely. The species with a PageRank index equal to 1 is the most important species according to the PageRank algorithm, ie, it is the most efficient nutrient catcher of the rooted food web 𝔽, the species with a PageRank index equal to 2 is the second most important, and so on.

Following Allesina and Pascual Allesina and Pascual, 2009], we are interested in the relative importance of each species in term of the support of other species. The corresponding species ranking can be obtained from the CheiRank vector P^*^ Chepelianskii, 2010, Zhirov et al., 2010]. Here, the elements of the corresponding stochastic matrix *S*^*^ are

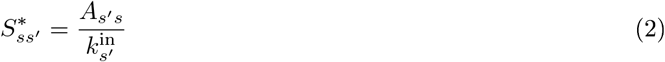

where 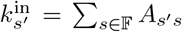 is the in-degree of the node *s′*, ie the number of preys of the species *s*′. The above defined out-degree 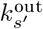 of the node *s*′ is then the number of predators of the species *s*′. Similarly to the PageRank algorithm, the steady state of the associated stochastic process is characterized by the CheiRank vector 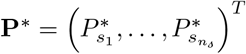 defined such as *S*^*^**P**^*^ = **P**^*^. Here, sorting the species by descending order of the value of their CheiRank probabilities, ie 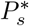 for the species *s*, allows to rank the species in accordance to their ability to spread nutrients through the food web and to support the other species Allesina and Pascual, 2009].

In Allesina and Pascual, 2009], Allesina and Pascual showed that this ranking gives a good extinction order to find *the most efficient route to collapse* of a trophic network. For each species *s* we assign a CheiRank index 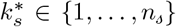 such as if 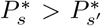, ie if the species *s*′ is a more efficient nutrient spreader than the species *s*′, then 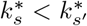, and conversely. The species with a CheiRank index equal to 1 is the most important species according to the CheiRank algorithm, i.e., the most efficient nutrient spreader, the species with a CheiRank index equal to 2 is the second most important, and so on.

### 1.2 PageRank algorithm for the landscape network

The nodes of the landscape network 𝕃 are the patches (or habitats) and the links are the paths between any couple of patches (see gray colored underlying graph in Fig. 1a). The number of patches is *n*_*p*_. We assume that the path from a node to another one can be taken either way, so the links are bidirectional and the landscape network is non directed. The associated stochastic matrix *S* has the elements

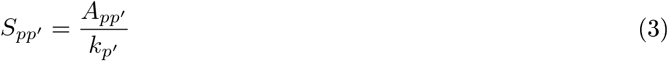

where *A*_*pp*_ ′ = 1 if there is a path between patches *p* and *p* ′, and *A*_*pp*_ ′ = 0 otherwise. The quantity *k*_*p*_ ′ = Σ_*p*∈_ *A*_*pp*_ ′ gives the number of patches which are connected to the patch *p*′. The stochastic matrix element *S*_*pp*_′ gives the rate of transition from the patch *p*′ to the patch *p*. We define then the PageRank vector 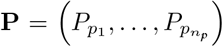 as the steady-state of the corresponding stochastic process, ***S*P** = **P**. The value of *P*_*p*_ is proportional to the number of times a random walker, forever wandering inside the landscape network 𝕃, hits the patch *p*. Similarly to the procedure followed previously in the section 1.1, we assign a PageRank index *K*_*p*_ to each patch. The patch with *K*_*p*_ = 1 is the most central patch according to the PageRank algorithm, ie, the most visited one, the one with *K*_*p*_ = 2 is the second most central, and so on. Otherwise stated, if *P*_*p*_ *> P*_*p*_ ′ or equivalently *K*_*p*_ *< K*_*p*_ ′, then the patch *p* is more often visited by a random walker than the patch *p* ′.

We note that the non directed nature of the landscape network 𝕃 would give a CheiRank probability vector exactly the same as the above defined PageRank probability vector since the inversion of the direction of the links leaves the landscape network 𝕃 unchanged.

### 1.3 CheiRank algorithm for the meta-food web

To a node of the meta-food web 𝕄 is attached a species *s* and a patch *p*. The elements of the adjacency matrix 𝒜 associated to the meta-food web 𝕄 are then

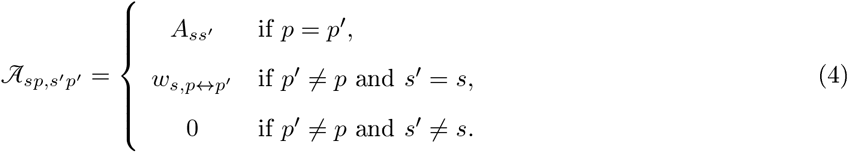

As long as we consider the same patch, i.e., *p* = *p*′, the elements 𝒜_*sp,s*′*p*_ of the meta-food web adjacency matrix 𝒜 are the same as those of the cloned rooted food web 𝔽, ie 𝒜_*sp,s′ p*_= *A*_*ss′*_ . We assume that the same rooted food web 𝔽 is duplicated on every patch. Once we consider different patches, ie *p* ≠ *p*′, a non zero value is possibly assigned to the same species adjacency matrix element 𝒜_*sp,sp*_′ = *w*_*s,p↔p*_′ if a link exists between patches *p* and *p*′, otherwise a zero value is assigned 𝒜_*sp,sp*_′ = 0. This representation allows coding for species-specific links in the landscape networks: not necessarily all species from a patch can disperse to the neighboring patches. Moreover, the non binary approach used here permits to use species specific strengths for dispersion links.

We assumed that no trophic interactions take place during the dispersion process. This implies that the energy transfer from one patch *p* to another patch *p*′ is always occurring between the same species within both patches. Hence, for two different species *s* and *s*′, and two different patches *p* and *p*′, 𝒜_*sp,s′ p*_′ = 0.

The inter-patch weight *w*_*s,p↔p*_ ′ is a priori dependent both on the considered species *s* and on the distance *d*_*p↔p*_ ′ between the patches *p* and *p* ′. It can be considered as the transition probability of the species *s* to cover the distance between the two patches *p* and *p* ′.

The CheiRank vector 𝒫^*^ of the meta-food web 𝕄 is obtained from the stochastic matrix 𝒮^*^ which the elements are

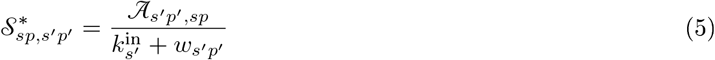

where 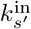 is the number of preys of the species *s*′ in the cloned rooted food web 𝔽 and *w*_*s′ p*_′ = Σ _*p*∈ 𝕃_ *w*_*s*′,*p↔p*_′ is the sum of the transit probabilities of the species *s*′ living in any patch *p* of the landscape network 𝕃 toward the patch *p*′.

As expected, the matrix 𝒮^*^ stochastic property is ensured as we have 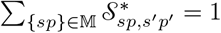. The CheiRank vector *𝒫*^*^ is defined such as *𝒮*^*^*𝒫*^*^ = *𝒫*^*^. The CheiRank probability 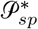 measures the ability of the species *s* living in the patch *p* to sustain all the other species living in the all the patches of the meta-food web 𝕄. The average CheiRank probability of the species *s* over the whole meta-food web is 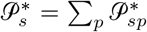, it measures the average ability of the species *s* to support the species living over the whole meta-food web 𝕄. The quantity 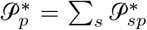 measures the ability of the patch *p* to sustain the entangled trophic network constituted by the meta-food web 𝕄.

As in the cases of the food web 𝔽 or of the landscape network 𝕃, it is possible to establish a ranking list 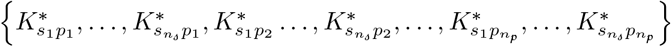, such as, if 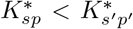, or equivalently 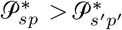, the species *s* in the patch *p* better supports the meta-community than the species *s*′ in the patch *p*_′_.

## 2 Case study on species importance: from local food webs to meta-communities

### 2.1 Methods

#### 2.1.1 Generation of the data

We generate random food webs with *n*_*𝓈*_ = 30 species using the niche model Williams and Martinez, 2000] with a connectance close to 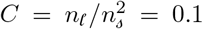. Here, *n*_*𝓁*_ is the number of trophic links in the food web. Fig 1b and c describe the transformation of a food web into the corresponding rooted food web 𝔽. Following Levine, 1980, Johnson et al., 2014], the trophic level TL_*s*_ of the species *s* is defined as 1 plus the average trophic level of its prey (the trophic level of basal species is set to 1). We associate a body mass *m*_*s*_ to species *s* depending on its trophic level TL_*s*_. The theoretical law giving the mass *m*_*s*_ of the species *s* according to its trophic level is 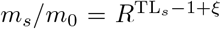 where *m*_0_ is the characteristic mass of the first trophic level, *R* is the average body-mass ratio between two species whose trophic levels differ by one unit, here we take *R* = 100, and *ξ* is a random variable sampled from a normal distribution with a mean of 0 and a standard deviation of 0.1.

The landscape network 𝕃 is a random geometric graph with *n*_*p*_ = 100 nodes. These nodes correspond to the patches of the landscape. The *n*_*p*_ patches are then randomly and homogeneously distributed in the unit square and two patches *p* and *p*_′_ are connected if they are separated by an Euclidean distance *d*_*p↔p* ′_ *< r*. Here, we take the threshold value *r* = 0.2. This graph is a non directed network. The species can transit between patches with no privileged direction. This landscape network is connected and its non directed nature ensures that there is no isolated node (or group of nodes) that would trap any random dynamics inside the network. In the following, a weight will be assigned to each link of the landscape network depending on the ability of a species to travel the corresponding distance between two patches.

We generate a meta-food web 𝕄 by replicating the same rooted food web 𝔽 in all the different landscape patches of the network 𝕃. We used 3 different scenarios to determine how species can disperse from a patch to another one and, consequently, to assign a weight *w*_*s,p↔p* ′_ to inter-patch links. In the first scenario, all species can disperse between two patches as soon as they are connected in the landscape network 𝕃. The second and third scenarios introduce a dispersion distance threshold that is species specific, and based on their body mass. More precisely:

- Type 1: All species are able to disperse between two patches as soon as they are connected in the landscape network. The inter-patch weight is species independent: *w*_*s,p↔p* ′_ = *w* for all the species. For this type of inter-patch weight, we consider the values *w* = 0.0001 corresponding to the regime where the duplicated food webs and the landscape network are almost totally decoupled (the nutrients are preferentially exchanged inside each patch and rarely travel from a patch to another), *w* = 0.1 corresponding to a moderate coupling, and *w* = 1 corresponding to a strong coupling for which the inter-patch weights are equal to the food web trophic links weights (the nutrients travel either from species to species or from patch to patch).
- Type 2: a species dependent weight *w*_*s,p↔p* ′_ which is equal to 1 if the species distance threshold *d*_*s*_ is greater than the inter-patch distance *d*_*p↔p* ′_ and equal to 0 otherwise,
- Type 3: the species dependent weight *w*_*s,p↔p* ′_ defined in (6) taking values inside the range [0, 1] depending on the distance *d*_*p↔p* ′_ between the two patches and on the species distance threshold *d*_*s*_.

Assuming that the most massive species, ie, the one with a mass *m*_max_ = max_*s*∈𝔽_*{m*_*s*_*}*, is able to reach a maximal distance of *r*, the normalized distance reachable by the species *s* is *d*_*s*_ = *r*(*m*_*s*_*/m*_max_)^*β*^ where *β* = 0.05 is the body mass scaling exponent. Hence, for the species *s*, the probability of transit 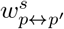 from the patch *p* to the patch *p*_′_, and *vice versa*, is assumed to follow a linear decrease with the actual distance *d*_*p↔p*′_ between the two patches

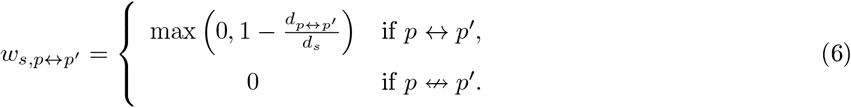

The first line in (6) concerns the case where a path actually exists between patches *p* and *p*_′_. If the maximum distance reachable by the species *s* is less than the distance between the patches *p* and *p*_′_, ie, *d*_*s*_ ≤ *d*_*p↔p* ′_, then the transition rate is zero, *w*_*s,p↔p*_′ = 0. In the opposite case, this rate *w*_*s,p↔p* ′_ = 1 − *d*_*p↔p* ′_ */d*_*s*_ decreases linearly with the inter-patch distance *d*_*p↔p* ′_ . The second line in (6) concerns the case when there is no geographical path between patches *p* and *p*_′_.

Mathematically, the meta-food web 𝕄 = 𝕃⊗𝔽, as depicted in Fig. 1a, is the tensor product of two networks, the landscape network 𝕃 and the rooted food web 𝔽. Hereafter, we will consider randomly generated hundreds of meta-food webs with (*n*_*𝓈*_ + 1) *× n*_*p*_ = 3100 nodes.

To assess how the position of a food web in the landscape network affects the importance of species locally, we examined how species’ CheiRank values change with the rank of patches from the landscape network.

#### 2.1.2 Dissimilarity of the food webs

We would like to compare the ranking of the species in the isolated food web 𝔽 with the relative ranking of the species inside a given patch *p* of the landscape 𝕃. For this purpose, we use the Kendall distance allowing to compare the differences between two ordered lists (in our case, two possible species rankings). Let us consider the *n*_*𝓈*_ species to which we assign the labels 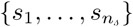. As introduced in section 1.1, the CheiRank algorithm allows to rank the species of the isolated food web 𝔽 according to their relative ability to support other species in the isolated food web: we obtain the CheiRank list 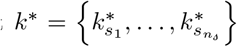. As introduced in section 1.3, the CheiRank algorithm allows to rank all pairs species-patch (*s, p*) of the meta-food web 𝕄. Focusing on a given patch *p*, we obtain the CheiRank list 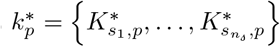. The Kendall distance Cicirello, 2020] between the CheiRank list *k*^*^ of the species in the isolated food web 𝔽 and the CheiRank list 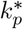 of the same species but living in the patch *p* of the meta-food web 𝕄 can be defined as

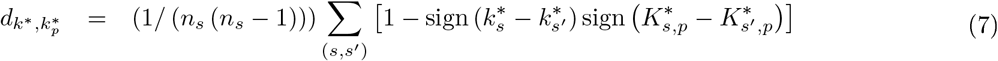

where the sum runs over all the different pairs (*s, s′*) of species, and the function sign (*x*) = *x/* |*x*| gives the sign of *x*, either +1 or −1. The Kendall distance 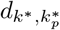 takes values from 0, for two lists *k*^*^ and 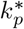 representing the same ordering of the *n*_*𝓈*_ species, to 1, for two lists that one represents the exact reverse ordering of the other, and conversely. The metric defined by (7) counts the number of pairwise disagreements between the two ranking lists *k*^*^ and 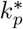.

### 2.2 Results

#### 2.2.1 Isolated food web versus meta-food web CheiRank analysis for an example food web

We present an illustration of how the importance of a species can change depending on the spatial context using an example of a rooted food web 𝔽 (Fig. 2a).

**Figure 2.**
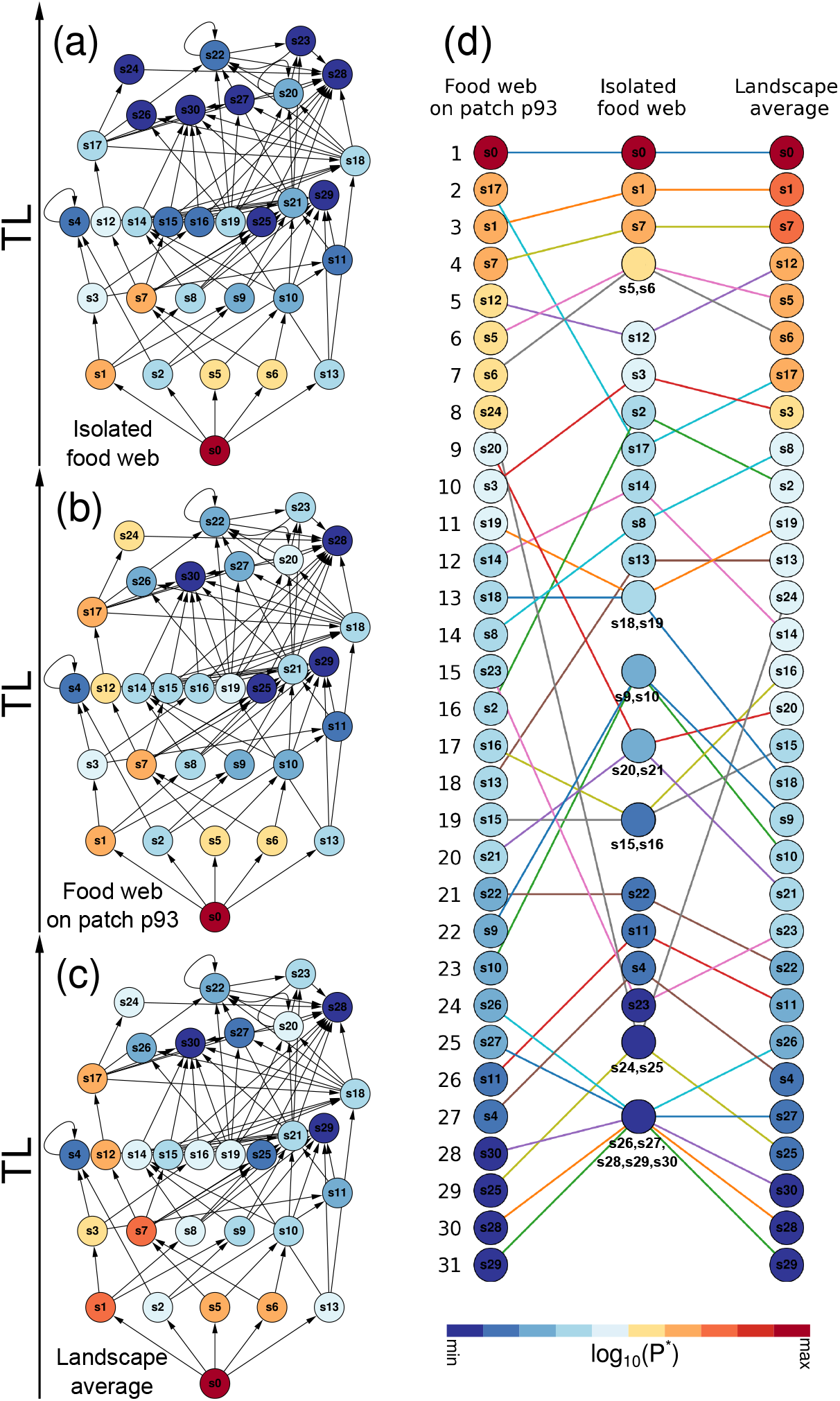
Typical food webs colored according to the CheiRank probabilities. The species *s* node is colored from dark blue (min) to dark red (max) according to its: **(a)** CheiRank probability value 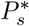 of the species in the isolated rooted food web 𝔽, **(b)** CheiRank probability value 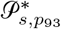 of the species in the patch *p*_93_ harboring the most dissimilar rooted food web to the isolated rooted food web 𝔽, **(c)** CheiRank probability value 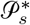 averaged over all the patches of the landscape network 𝕃. The minimum and the maximum values of the CheiRank probabilities are : (a) 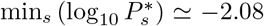 and 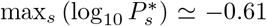, (b) 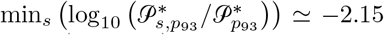 and 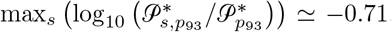, (c) 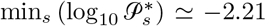 and 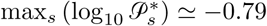. For the sake of visibility, the recycling links “species → root” (as an example, see red colored dotted links in Fig. 1c) are not shown. The panel **(d)** shows the ranking of the species: (middle column) living in the isolated food web 𝔽, (left column) living on the patch *p*_93_, (right column) averaged over all the patches of the landscape network 𝕃. For the isolated food web, the corresponding minimum rank is assigned to tied species.

Trivially, the root node *s*_0_ has by far the highest CheiRank probability for all food webs, the isolated one (Fig. 2a), but also all food webs in the meta-community context (Fig. 2bc). This is obvious since it supports the whole food web and its removal would mechanically leads to the extinction of all the other species. Then, species considered as important tend to be located in the lower part of the isolated rooted food web 𝔽 (Fig. 2a), but with important differences within trophic levels (e.g., basal species *s*_1_, *s*_5_, *s*_6_, *s*_7_ from the isolated food web have CheiRank probabilities greater than the ones of higher trophic levels. Indeed, according to the CheiRank algorithm, the more a species supports important supporting species, the more it is an important supporting species (recurrence definition).

We duplicated now the rooted food web 𝔽 into each patch of the landscape network 𝕃 in order to construct the meta-food web 𝕄. We use a species transit probability that is dependent of species body mass, as above defined in the type 3 scenario (6). We will begin by outlining the differences in CheiRank distributions between the isolated food web 𝔽 (Fig. 2a) and its counterpart in the meta-community context which exhibits less similarity to it (Fig. 2b, the most dissimilar patch being here *p*_93_). In a second step we will compare the CheiRank distributions of the isolated rooted food web to an averaged CheiRank distribution accross all patches (Fig. 2c).

We observe, within the patch *p*_93_ (Fig. 2b), that the species with the highest CheiRank probabilities are less systematically located at the bottom of the sub-network (species with low TL). This difference with the isolated food web 𝔽 (Fig. 2a) is due to possible displacements of species through the whole landscape 𝕃. From the species ranking (Fig. 2d), we observe that species *s*_17_, with rank 9 in the isolated food web, is the most important living species in the patch *p*_93_, with rank 2 (the non living root node has the rank 1). Also species *s*_24_ gains 17 places since it passes from rank 25 in the isolated food web to rank 8 in the patch *p*_93_. Additional species obtain a significantly better ranking in patch *p*_93_ (e.g., species *s*_20_ and *s*_23_) whereas others obtain a significantly lesser ranking (e.g., TL=1 species *s*_3_ and *s*_13_ which passes from rank 7 to 10 and from rank 12 to 18, respectively). Hence, from one patch to another, the relative ability of a species to support the others can substantially change.

Also, from Fig. 2d, we observe that the existing ties in the isolated food web 𝔽 ranking are removed due to the different species displacement capabilities over the landscape 𝕃 (e.g., the species *s*_5_ and *s*_6_, which are equivalent for the CheiRank algorithm as they both provide food for species *s*_7_ and *s*_10_, have the same ranking in the isolated food web and different rankings in the patch *p*_93_).

To obtain the average CheiRank probability distribution 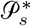 for the meta-community in Fig. 2c we summed the CheiRank probabilities 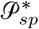 over all the patches (see section 1.3). The obtained space averaged CheiRank probability distribution clearly exhibits differences with the isolated rooted food web 𝔽 (Fig. 2a). In particular, we observe a preferential food path

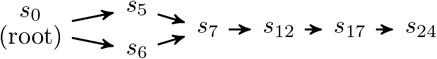

which, on average, sustains the whole meta-food web 𝕄 (and that is also visible on *p*_93_). In addition, some top predators, e.g., *s*_20_, *s*_23_ and *s*_24_, have a better averaged CheiRank probability in the meta-food web 𝕄 (see Fig. 2c) than in the isolated rooted food web 𝔽 (see Fig. 2a). These top predators participate more in the sustainment of the meta-food web than in the isolated rooted food web. Indeed, the mobility, increasing with species’ body-masses, favors the spreading of top predators over the landscape which can be in return sustained by the preys living on all the different patches. Particularly, from Fig. 2d (right column), although the species *s*_24_ (rank 13) is a top predator with TL>4 (see Fig. 2a), it plays a non-negligible role in maintaining the energetic integrity of the meta-community.

#### 2.2.2 Statistical analysis of the dissimilarity between an isolated food web and the meta-food web

In the following we assess how this consideration of the landscape network alters our definition of important species using a replicated approach on the landscape. We consider the 3 different types of inter-patch weights *w*_*s,p*↔*p′*_ ensuring the diffusion of the species from patch to patch over the landscape: 1) independent of species, 2) different links depending on species body mass, same weight for all species and 3) species specific link and link weight (see section 2.1.1 for a detailed description).

As before, we consider a meta-food web 𝕄 constituted by an underlying landscape 𝕃 of patches hosting in each patch the same rooted food web 𝔽. We however created 100 meta-food webs 𝕄 that differ by the rooted food web used (100 different food webs were generated and combined with the same Landscape network). This procedure is followed for the three scenarios of species mobility. For each generated meta-food web, we compute the CheiRank vector and we rank all the species living in the landscape according to their CheiRank probability.

For a given meta-food web 𝕄, we compute the Kendall distances 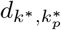 (7) between the CheiRank list of the species belonging to the isolated food web 𝔽 and the CheiRank list of the species living inside each patch of the landscape hosting all the same food web 𝔽. In Fig. 3, we present, for each landscape patch, the average of the Kendall distance over the 100 randomly generated meta-food webs.

**Figure 3.**
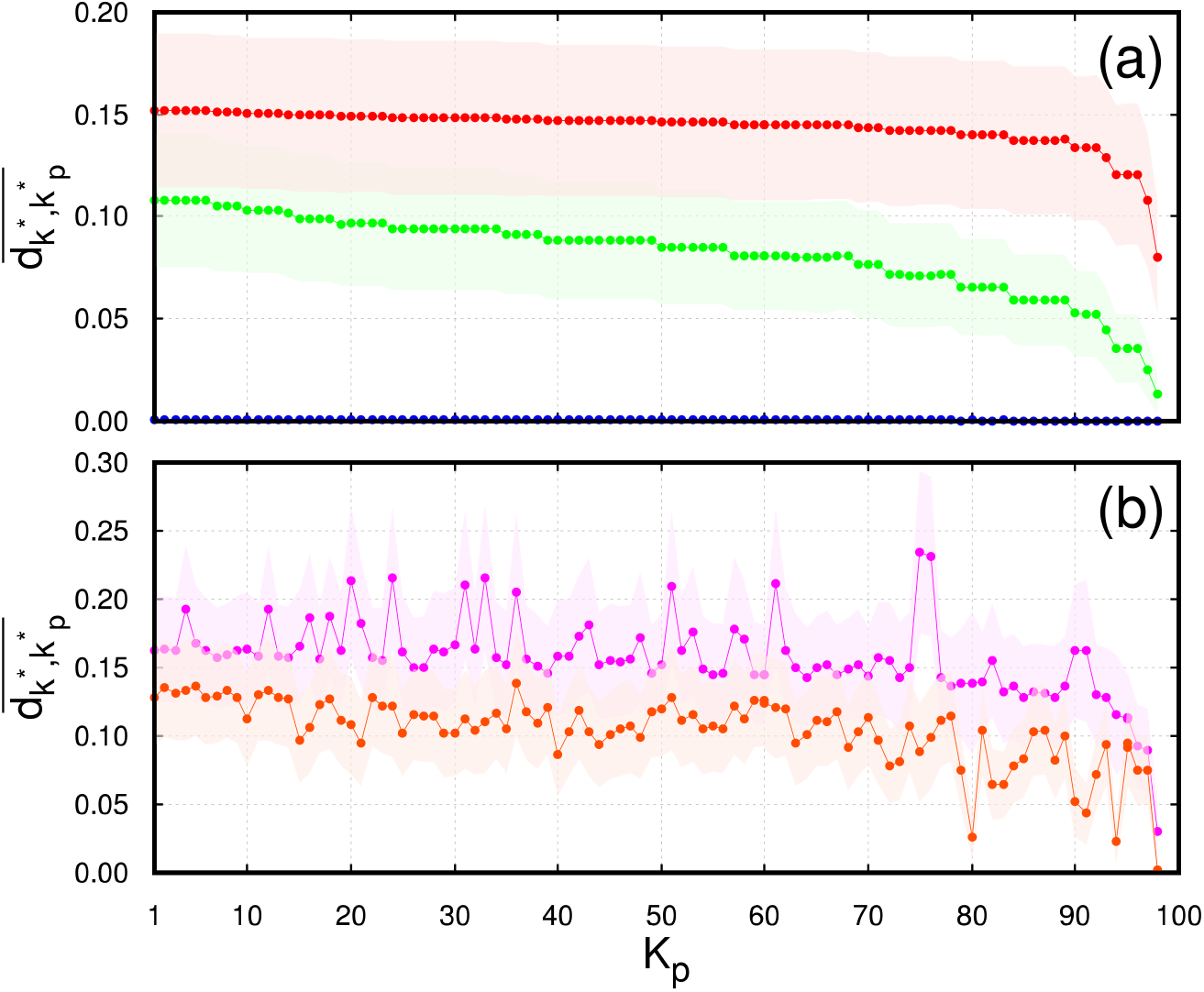
Dissimilarity between the food webs located on the patches and the isolated food web. The Kendall distance 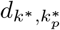 is computed between the CheiRank list of species in the isolated rooted food web 𝔽 and the CheiRank list of species in the food web located on a given patch of the meta-food web 𝕄. The average 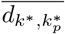 is done over 100 randomly generated meta-food webs keeping the same underlying landscape network and randomly generating new cloned rooted food web 𝔽. Along the abscissa, the patches are ordered by their PageRank indices *K* computed from the non directed landscape network 𝕃. **Panel (a)** present the computed distances for the type 1 species independent inter-patch weights with *w* = 0.0001 (blue), *w* = 0.01 (green), and *w* = 1 (red). The minimum, **the mean**, the standard deviation, and the maximum of the averaged Kendall distance 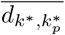 are 0.0001, **0.0004**, 0.0001, 0.0005 for *w* = 0.0001, 0.0134, **0.0817**, 0.0196, 0.1079 for *w* = 0.01, and 0.0801, **0.1437**, 0.01, 0.1520 for *w* = 1, **Panel (b)** present the computed distances for the species dependent inter-patch weights, type 2 (magenta) and type 3 (orange). The minimum, **the mean**, the standard deviation, and the maximum of the averaged Kendall distance 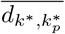 are 0.0305, **0.1569**, 0.0284, 0.2346 for type 2 and 0.0025, **0.1045**, 0.0243, 0.1383 for type 3. The shaded areas surrounding the different points delimit the ± σ standard deviation range associated to the random food webs distribution.

The panel Fig. 3a shows for each patch *p* the average 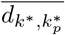 of the Kendall distance over the 100 random realizations of the meta-food web and for the species independent inter-patch weights (type 1). In abscissa, the landscape patches are ordered according to their respective PageRank index *K*_*p*_ obtained from the PageRank vector associated to the landscape network 𝕃. As expected, the almost decoupled regime *w* = 0.0001 (blue points in Fig. 3a) leads to a Kendall distance close to 0 for all the patches. For the moderate coupling regime *w* = 0.1 (green points in Fig. 3a), the distance 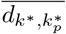 globally drops as the landscape PageRank index *K*_*p*_ of the patch increases. Indeed, we expect that the species living on the most (less) central patches are the most (less) impacted by the structure of the underlying landscape network. In the strong coupling regime *w* = 1 (red points in Fig. 3a), the drop is less pronounced. In this regime, the weights of the food web links and the inter-patch weights are equal. Consequently, we can consider that the duplicated food webs and the landscape network are totally merged into a single whole network. Otherwise stated, nutrients can pass as easily from one species to another as from one patch to another.

The panel Fig. 3b presents the average Kendall distance 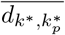 for the species dependent inter-patch weights (type 2 and 3). This types of species diffusion are by far more realistic as they take account of the body mass dependent ability of each species *s* to travel over a given distance (i.e, the distance threshold *d*_*s*_ is body mass specific). Globally, as observed in Fig. 3a for the species independent type of inter-patch weights, the curves follow the same trend, ie, the food webs located on the most central patches of the landscape have the largest Kendall distance from the isolated food web, and conversely, the less central ones have the smallest distance. But, more interestingly, the drop of the Kendall distance with the landscape PageRank centrality of the patches is only a trend as the drop is no more monotonous and non negligible fluctuations of the mean now appear: two patches with comparable PageRank indices *K*_*p*_ may exhibit an up to 0.1 difference of degrees of dissimilarity with the isolated rooted food web. The fluctuations are more pronounced for the type 2 species diffusion (magenta points) than for the type 3 species diffusion (orange points) which is more realistic. Indeed, even not very central patches in the landscape according to the PageRank algorithm harbor food webs with the maximum dissimilarity with the isolated rooted food web. The top PageRank patches are not necessarily the most dissimilar ones with the genuine food web; this is particularly the case for the type 2 species mobility model. For this type two mobility model, we can see that the distribution of species importance in the rooted food webs from patches with PageRank indices 75 and 76 are the one that differ the most from the isolated food web. For the Type 3 mobility scenario, the strongest difference is observed for the patch with a PageRank index of 36 .

The most important feature is, independently of the species mobility model, that the food web’s CheiRank can present in average up to 15% differences with the genuine food web if we consider its environment and the possibility for species to travel throughout the landscape.

Based on these results, we ran a subsequent analysis to assess how the position of a food web in the landscape network affects the importance of species locally. Using the same procedure as described before, we generated 100 different landscape networks (for which nodes were linked using the type 3 scenario), each associated to 100 different food webs. We observe (Fig. 4) a decrease in species importance depending on their position on the landscape networks. This result therefore indicates that the most important species in meta-communities will mostly be observed in landscape patches considered as important.

**Figure 4.**
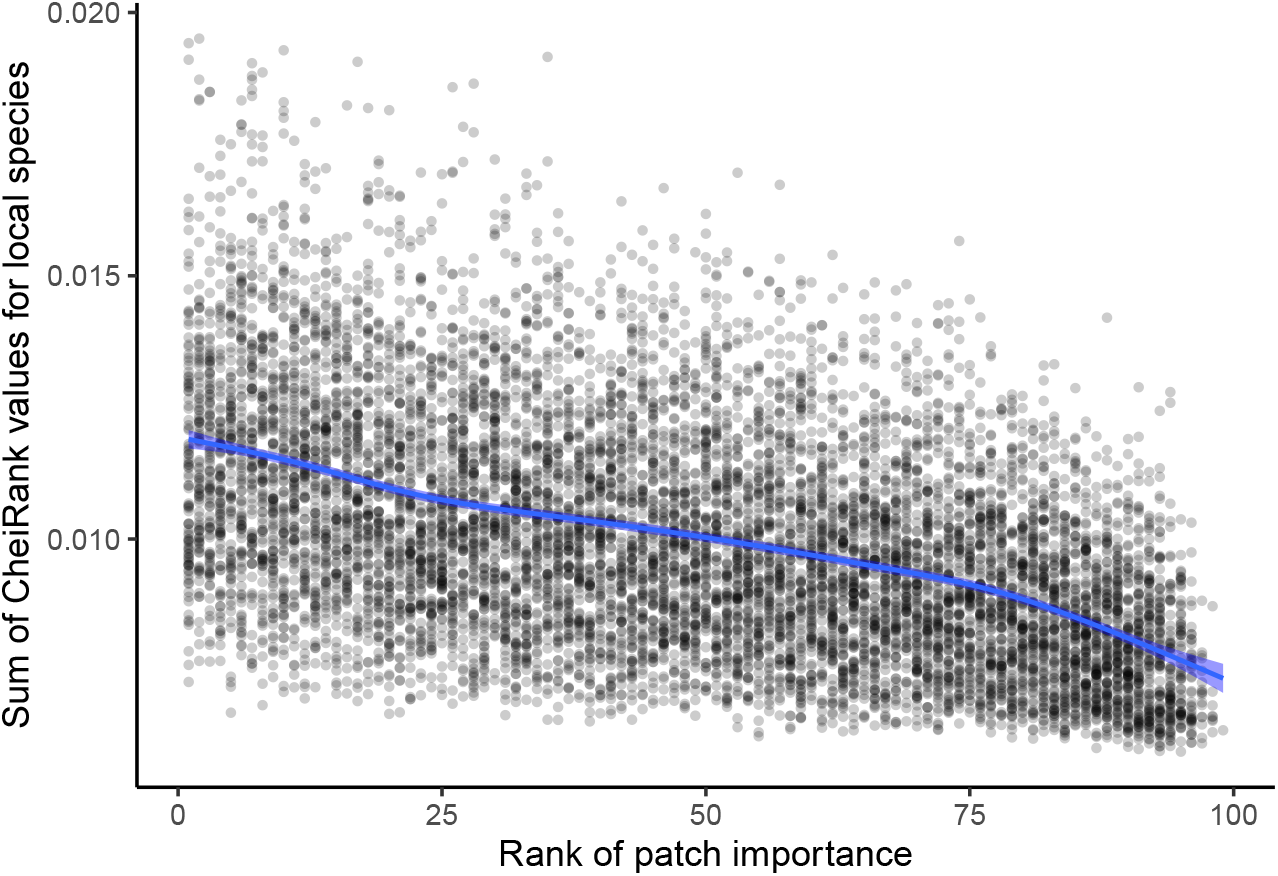
Effect of patch importance on local species CheiRank values. Each dot represents the sum of CheiRank values of species on a patch of a landscape network, average over 100 different food webs. The blue line represents predictions from a GAM model using patch rank as a predictor variable. The blue shaded area represents the 95% confidence interval on model predictions.

## 3 Discussion

By proposing a method to estimate species’ importance in meta-food webs, we were able to draw two main conclusions. First, species importance between the isolated food web and the different local food webs in the community changes depending on our dispersion scenario (i.e. all species have the same dispersion abilities vs species dependent abilities). When considering more realistic dispersion models for species (species-dependent inter-patch weights), fluctuations of dissimilarity no longer follow a monotonic pattern. As the model used here only relates to the topology of the two networks -the food web and the landscape network -this fluctuation is a signature of the entanglement of the food webs and the underlying landscape network through the body-mass-dependent species mobility. This means that the importance of a species locally will relate to both its position in the food web Allesina and Pascual, 2009, Jordán, 2009] and its capacity to move across patches in a landscape. Given the recent advocacy of functional trait approaches for identifying key species in ecosystems ( Schleuning et al., 2023, Brun et al., 2022]), our findings underscore the significance of encompassing not only traits associated with the roles and trophic positions of species within their communities, but also traits linked to dispersion.

Second, we show the importance of considering the local context and the positioning of communities within their global environment when identifying important species. Indeed, we found a substantial variation of species importance, both between food webs located in different patches of a meta-food web and between isolated food webs and food webs embedded in space. Overall, food webs from central patches are the ones that differ the most from the isolated food webs, meaning that what are the important species in these central patches can be di cult to predict from the isolated network. As we observed that species from these central patches tend to be the most important in meta-communities, our results stress the need for a global consideration of space in the identification of important species. This context dependency of the rank of species importance raises a fundamental conclusion: species importance is not necessarily a species characteristic associated with its identity, but a property that varies depending on the ecological context: a species considered as the most important in a patch of the landscape network might not be in another patch. On the whole, it means that the importance of a species is influenced by both intrinsic factors (dispersal capacity, trophic position) and extrinsic factors (position of the patch in the landscape network).

For the sake of completeness, we have also computed the dissimilarity in terms of the PageRank centrality (not shown here) instead of the CheiRank centrality. While the CheiRank centrality quantifies the ability of species to distribute energy in the food web or the meta-community the PageRank centrality quantifies its ability to catch energy. The above conclusions drawn for the CheiRank centrality (see Figs. 3ab) stand also for the PageRank centrality (see Fig. SI I). Consequently, on average, the patches which capture the most e ciently the nutrients are also the ones disseminating the most. These patches therefore act as energetic hubs of the meta-food web. We note that such a combined CheiRank-PageRank analysis has been already applied to various directed networks such as the international trade network (see e.g., Coquidé et al., 2020]), protein-protein interaction networks (see e.g., Lages et al., 2018]), and WWW-like networks (see e.g., Zhirov et al., 2010, Rollin et al., 2019]). We leave such study of meta-food webs with underlying directed landscape network for a further paper.

Although each duplicated food web is directed, the non directed nature of the underlying landscape network dominates the meta-food web. It would be interesting to investigate the case of a directed landscape network modeling possible one-way inter-patch paths, dead ends, or even asymmetric access to some habitat (for instance because of relief in mountainous areas). We argue that in this case the combined CheiRank-PageRank analysis of the corresponding meta-food web will permit to better integrate habitat characteristic into model’s predictions. Spatial links could for instance be parametrised based on the energy cost of dispersion Berti et al., 2022] to further integrate habitat complexity as well as ecological and physiological processes.

Nowadays, the scientific community stresses the need of protecting native and endangered biodiversity and preserve ecosystems, by among other restoring or preserving habitat connectivity Cushman et al., 2013] and giving a special care to important species Valls et al., 2015]. Our model is a first attempt to bridge these two approaches by potentially allowing the identification of which species in which patches are important to preserve in order to maintain the integrity of energy transfer in a landscape. While we must acknowledge that current applications are yet limited by the amount of information needed to build meta-networks, the use of species functional traits offers some promising results. Body mass is currently utilised with success to predict the distribution of trophic interactions among species Gravel et al., 2013], and its predictive power could further be enhanced by incorporating additional traits such as metabolic type or foraging behavior Li et al., 2023]. Regarding species dispersal ability, body mass also appears to be a fundamental trait Hirt et al., 2018] that can be coupled with movement mode (i.e. crawling flying, …). Last, species occurrence in the different patches can be estimated with models based on habitat suitability Elith and Leathwick, 2009]. As such, the method we present has a strong potential for synergies with the recent development of functional trait approaches for conservation ecology Gallagher et al., 2021].

## Supplementary Information I

**Figure SI I:**
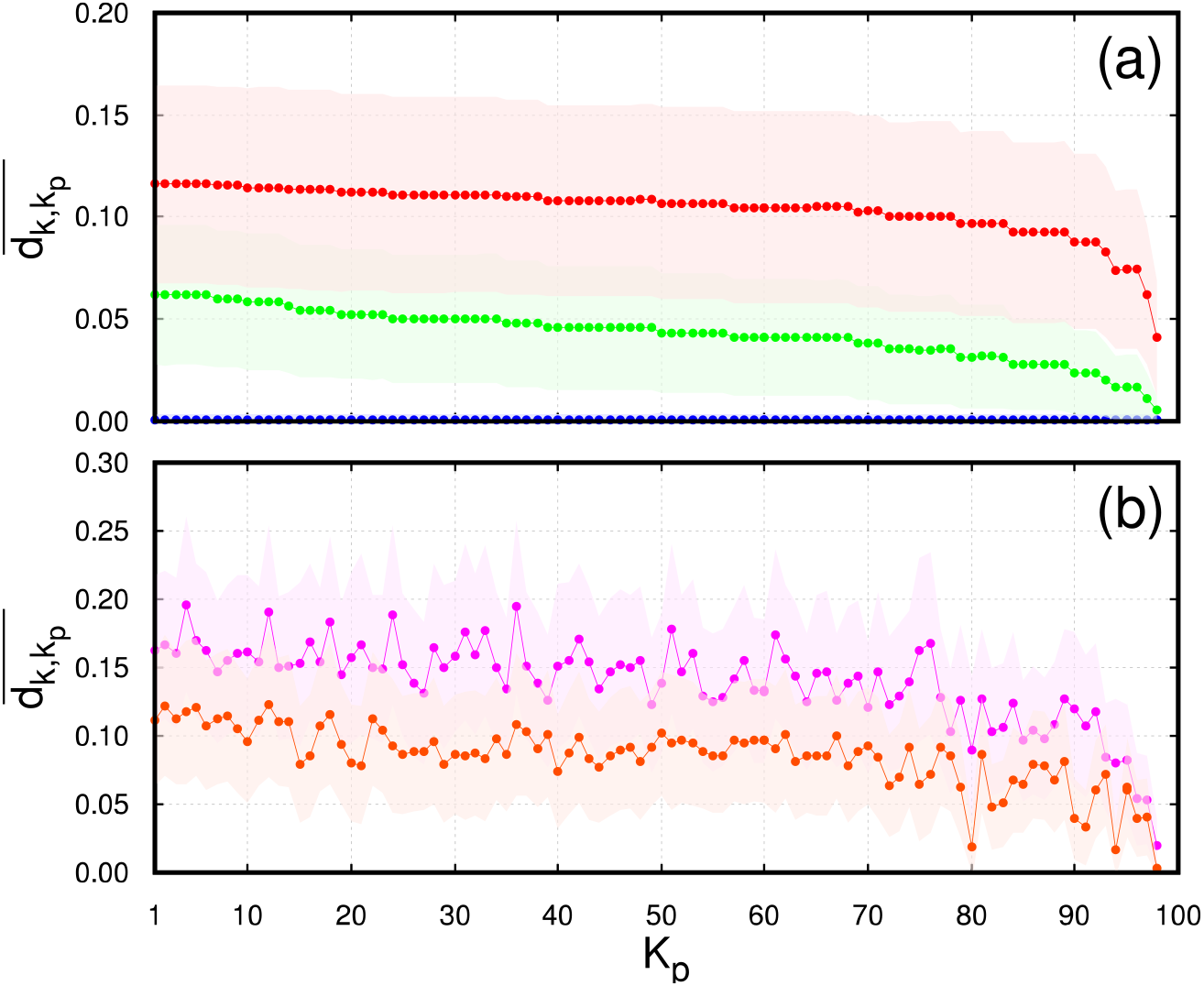
Dissimilarity between the food webs located on the patches and the isolated food web. The Kendall distance 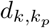 is computed between the PageRank list of species in the isolated rooted food web 𝕄 and the PageRank list of species in the food web located on a given patch of the meta-food web 𝕄. The average 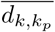 is done over 100 randomly generated meta-food webs keeping the same underlying landscape network and randomly generating new cloned rooted food web 𝔽. Along the abscissa, the patches are ordered by their PageRank indices *K* computed from the non directed landscape network 𝕃. **Panel (a)** present the computed distances for the type 1 species independent inter-patch weights with *w* = 0.0001 (blue), *w* = 0.01 (green), and *w* = 1 (red). The minimum, **the mean**, the standard deviation, and the maximum of the averaged Kendall distance 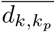 are 0.0006, **0.0007**, 4*×*10^−5^, 0.0007 for *w* = 0.0001, 0.0053, **0.0427**, 0.0123, 0.0622 for *w* = 0.01, and 0.0408, **0.1039**, 0.0122, 0.1160 for *w* = 1, **Panel (b)** present the computed distances for the species dependent inter-patch weights, type 2 (magenta) and type 3 (orange). The minimum, the mean, the standard deviation, and the maximum of the averaged Kendall distance 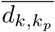 are 0.0194, **0.1393**, 0.0303, 0.1962 for type 2 and 0.0028, **0.0853**, 0.0224, 0.1232 for type 3. The shaded areas surrounding the different points delimit the ±σ standard deviation range associated to the random food webs distribution.

